# Life Inside a Dinosaur Bone: a Thriving Microbiome

**DOI:** 10.1101/400176

**Authors:** Evan T. Saitta, Renxing Liang, Chui Y. Lau, Caleb M. Brown, Nicholas R. Longrich, Thomas G. Kaye, Ben J. Novak, Steven Salzberg, Paul Donohoe, Marc Dickinson, Jakob Vinther, Ian D. Bull, Richard A. Brooker, Peter Martin, Geoffrey D. Abbott, Timothy D. J. Knowles, Kirsty Penkman, Tullis C. Onstott

## Abstract

Fossils were long thought to lack original organic material, but the discovery of organic molecules in fossils and sub-fossils, thousands to millions of years old, has demonstrated the potential of fossil organics to provide radical new insights into the fossil record. How long different organics can persist remains unclear, however. Non-avian dinosaur bone has been hypothesised to preserve endogenous organics including collagen, osteocytes, and blood vessels, but proteins and labile lipids are unstable during diagenesis or over long periods of time. Furthermore, bone is porous and an open system, allowing microbial and organic flux. Some of these organics within fossil bone have therefore been identified as either contamination or microbial biofilm, rather than original organics. Here, we use biological and chemical analyses of Late Cretaceous dinosaur bones and sediment matrix to show that dinosaur bone hosts a diverse microbiome. Fossils and matrix were freshly-excavated, aseptically-acquired, and then analysed using microscopy, spectroscopy, chromatography, spectrometry, DNA extraction, and 16S rRNA amplicon sequencing. The fossil organics differ from modern bone collagen chemically and structurally. A key finding is that 16S rRNA amplicon sequencing reveals that the subterranean fossil bones host a unique, living microbiome distinct from that of the surrounding sediment. Even in the subsurface, dinosaur bone is biologically active and behaves as an open system, attracting microbes that might alter original organics or complicate the identification of original organics. These results suggest caution regarding claims of dinosaur bone ‘soft tissue’ preservation and illustrate a potential role for microbial communities in post-burial taphonomy.

## Introduction

Fossils have often been thought to contain little original organic material due to decay and diagenesis. However, discoveries of ancient DNA (Orlando *et al.* 2013) and peptide (Demarchi *et al.* 2016) sequences in sub-fossils as well as very ancient biomolecules such as sterols (Melendez *et al.* 2013) and melanin (Vinther *et al.* 2008) in fossils challenge this view. These finds show that organic remains can potentially persist for thousands to even millions of years depending on the biomolecule and environmental conditions. These organic molecular fossils can potentially shed light on the biology and evolution of extinct organisms (e.g., colour, morphology, behaviour, phylogeny), providing unique insights into past life and the origins of present life. In theory, organic remains capable of surviving millions or tens of millions of years may offer palaeontologists new insights into the biology of organisms distantly related to any living species, such as dinosaurs. However, exactly how long different types of organic molecules can potentially survive remains debated.

Dinosaur bone has been reported to contain endogenous organics such as collagen, osteocytes, and blood vessels. These reports, if verified, could radically change the study of macroevolution and the physiology of extinct organisms given the immense potential of protein sequence data to shed light on the biology and systematics of extinct organisms (Pawlicki *et al.* 1966; Schweitzer *et al.* 2005a, 2005b, 2007a, 2007b, 2008, 2009, 2013, 2014, 2016; Asara *et al.* 2007; Organ *et al.* 2008; Schweitzer 2011; Bertazzo *et al.* 2015; Cleland *et al.* 2015; Schroeter *et al.* 2017). Most of these reports rely on structural observations, mass spectrometry, and immunohistochemistry. It makes sense to search for collagen in ancient remains since such sub-fossils and fossils are often bone, dentine, or enamel and these calcified tissues have protein compositions dominated by collagen, which relative to many other vertebrate proteins is robust due to its decay resistance, triple helical quaternary structure, and high concentration of thermally stable amino acids (Engel & Bächinger 2005; Persikov *et al.* 2005; Sansom *et al.* 2010; Wang *et al.* 2012). However, others have criticised such mass spectral data, saying that it represents laboratory or environmental contamination (Buckley *et al.* 2008, 2017; Bern *et al.* 2009) or statistical artefacts (Pevzner *et al.* 2008). Furthermore, antibodies are known to cause occasional false positives (True 2008). Organics within fossil bone of appreciable age/thermal maturity producing ‘vessel’- and ‘cell’-shaped molds have alternatively been identified as biofilm (Kaye *et al.* 2008).

Such structures, if indeed Mesozoic ‘soft tissue’, would be expected to consist primarily of extracellular structural proteins and phospholipids of cell membranes, which are unstable through diagenesis and deep time (Bada 1998; Briggs & Summons 2014). While protein sequences are lost through hydrolysis of peptide bonds, phospholipids hydrolyse at their ester bonds, freeing fatty acids from glycerol-phosphate polar heads (Eglinton & Logan 1991; Zuidam & Crommelin 1995). The resulting free fatty acids, however, can polymerise *in situ* to form kerogen-like aliphatic hydrocarbons which are stable through diagenesis (Stankiewicz *et al.* 2000; Gupta *et al.* 2006a, 2006b, 2007a, 2007b, 2008, 2009). The conversion of the hydrocarbon tails of phospholipids into kerogen is far more likely than preserving a protein sequence of amino acids. One survey of amino acids in fossil bone (Armstrong *et al.* 1983) yielded data in which fossils older than the Upper Pleistocene no longer had amino acid compositions similar to fresh bone proteins. Additionally, amino acid concentration decreased from recent bone to Mesozoic bone, while racemisation increased sharply from recent bone to Upper Pleistocene bone, but then gradually decreased from that point through to Mesozoic bone, suggesting protein loss and contamination (see supplemental material for a reanalysis of this data from Armstrong *et al.* 1983). However, partially intact Pliocene peptides about 3.4 Ma have been verified from exceptionally cold environments (Rybczynski *et al.* 2013) and under what, for now at least, seems like unique molecular preservational mechanisms in the calcite crystals of eggshell from 3.8 Ma (Demarchi *et al.* 2016). Both of these examples, however, a far younger than Mesozoic fossils.

Even simple estimations do not predict protein survival in the deep geologic record. Assuming fairly average human body composition (Janaway *et al.* 2009) and mass, it only takes ∼5 % of the water already present in the body to hydrolyse all of the peptide bonds in the proteome (supplemental material). This calculation assumes a closed system with no endogenous water, and it seems unlikely that any fossil matrix would be anhydrous throughout its entire taphonomic history. For example, it requires a significant amount of diagenetic alteration to fossilise resin into desiccated copal and amber (Langenheim 1969, 1990; Lambert & Frye 1982; Mills *et al.* 1984; Pike 1993; Ragazzi *et al.* 1993; Villanueva-García *et al.* 2005). Calculating exponential decay curves assuming first order kinetics paints an even more pessimistic picture. Modifications at terminal regions and internal peptide bonds shielded by steric effects can result in longer observed half lives of peptide bonds under hydrolysis (Kahne & Still 1988; Radzicka *et al.* 1996; Testa & Mayer 2003). At 25 °C and neutral pH, peptide bond half lives as a result of uncatalysed hydrolysis for the relatively stable acetylglycylglycine (C-terminal), acetylglycylglycine *N*-methylamide (internal), and the dipeptide glycylglycine are 500, 600, and 350 years, respectively (Radzicka *et al.* 1996). Even assuming a very conservative half life of 600 years for all peptide bonds in the average human body at arguably unextreme conditions (25 °C and neutral pH), no bonds would remain after ∼51,487 years (supplemental material):

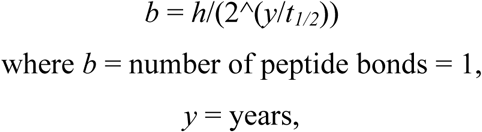

*h* = estimated number of peptide bonds in average human body = 6.78844 × 10^25, and *t1/2* = half life in years under uncatalysed hydrolysis = 600.

The half life in this case is 3 orders of magnitude too short in order to get peptide bonds surviving into the Mesozoic (∼66 Ma); a half life of ∼769,130 years would be required. This does not even take into consideration environmental or diagenetic increases in temperature or pH fluctuations, nor does it take into account scavenging or microbial/autolytic decay. Of course, these values are based on some very extreme assumptions and should not be taken as precise estimates, but rather, as framing the enormity of the challenge for Mesozoic protein survival. Empirically derived estimates for collagen and osteocalcin upper age limits based on experimentally observed gelatinisation and Gla-rich mid-region epitope loss, respectively, can give widely different estimates at 20 °C: 15,000 years for collagen and 580,000 years for osteocalcin. The estimates vary according to temperature. For example, at 0 °C, the upper age limit for collagen and osteocalcin are estimated at 2,700,000 and 110,000,000 years, respectively (Nielsen-Marsh 2002). Even frozen collagen by these estimates fails to survive long enough for the possibility of survival in Mesozoic specimens, and no Mesozoic fossils have been preserved frozen since they predate the appearance of the current polar ice caps.

The kinetics of thermal instability under non-enzymatic reactions are just one hurdle that such ‘soft tissues’ would have to clear. Bone is also an open system (Bada *et al.* 1999), allowing for organic and microbial influx. Invasion of microbes into the bone could lead to the enzymatic degradation of endogenous organics (in addition to any autolytic degradation from endogenous enzymes) and mobile breakdown products of organics can be lost from the bone into the surroundings.

Here, chemical and biological analyses of freshly-collected, aseptically-acquired, Late Cretaceous surface-eroded and excavated subterranean dinosaur bones, compared to associated sediment and soil, younger fossil, and modern bone controls, show evidence for a microbiome. Analyses were conducted using variable pressure scanning electron microscopy (VPSEM), energy dispersive X-ray spectroscopy (EDS), light microscopy, attenuated total reflectance Fourier-transform infrared spectroscopy (ATR FTIR), pyrolysis-gas chromatography-mass spectrometry (Py-GC-MS), high performance liquid chromatography (HPLC), radiocarbon accelerator mass spectrometry (AMS), Qubit fluorometer, epifluorescence microscopy (propidium iodide (PI) and SYTO 9 staining), and 16S rRNA amplicon sequencing.

In addition to finding little evidence for the preservation of original proteinaceous compounds, our findings suggest that bones not only act as open systems just after death and exhumation, but also as fossils in the subsurface. Microbial communities appear to be thriving inside the dinosaur bones collected here.

## Materials and Methods

### Aseptic fossil acquisition

Samples of Late Cretaceous fossil dinosaur bone, along with associated sediment and soil controls were obtained from the Dinosaur Park Formation (Late Campanian) in Dinosaur Provincial Park, Alberta, Canada (supplemental material). The Dinosaur Park Formation is a well-sampled alluvial-paralic unit deposited during a transgressive phase of the Western Interior Seaway, for which a diverse vertebrate fauna has been documented over more than a century of collection (Currie and Koppelhus, 2005). The bone samples were collected from a monodominant bonebed (BB180) of the centrosaurine *Centrosaurus apertus* (Ornithischia; Ceratopsidae), located 3 m above the contact with the underlying Oldman Formation (precise location data available at the Royal Tyrrell Museum of Palaeontology). The mudstone-hosted bone-bearing horizon is an aggregation of disarticulated but densely packed bones, with a vertical relief of 15–20 cm. Similar to other ceratopsid bonebeds from the same stratigraphic interval (Ryan *et al*. 2001; Eberth and Getty, 2005), the recovered skeletal remains are nearly exclusively from Ceratopsidae, and with all diagnostic ceratopsid material assignable to *Centrosaurus apertus*, with the site interpreted as a mass-death assemblage. Fossil material was collected under a Park Research and Collection Permit (No. 16-101) from Alberta Tourism, Parks and Recreation, as well as a Permit to Excavate Palaeontological Resources (No. 16-026) from Alberta Culture and Tourism and the Royal Tyrrell Museum of Palaeontology, both issued to C. M. Brown.

Sandstone and mudstone overburden was removed with pick axe and shovel (∼1 m into the hill and ∼1 m deep) to make accessible a previously unexcavated region of the bonebed, stopping within ∼10 cm of the known bone-bearing horizon. A few hours after commencement of overburden removal, excavation of the mudstone to the bone-bearing horizon was conducted using awl and scalpel. Subterranean *Centrosaurus* bones (identified as a small rib and a tibia) were first discovered non-aseptically, exposing them to the air in a manner identical to typical palaeontological excavation and allowing for rapid detection of bones.

At this point, aseptic techniques were then implemented to expose more of the bone in order to determine its size and orientation. It is worth qualifying the usage of the term ‘aseptic’ in this study. Paleontological field techniques have changed little over the last century, and it is likely near impossible to excavate fossils in an absolutely sterile manner (e.g., the process of matrix removal induces exposure, the wind can carry environmental contaminants onto exposed fossils, etc.). In light of this, the term ‘aseptic’ is used here to to acknowledge the inability to provide completely sterile sampling conditions, while still indicating that efforts are taken to reduce introducing contamination to the samples to an extent that might influence results. Our success at reasonably reducing contamination is evidenced by the fact that our samples yielded consistent and interpretable results.

During aseptic excavation and sampling, nitrile gloves washed in 70 % ethanol and a facemask were worn. All tools (i.e., awl, scalpel, dremel saw) were sterilised with 10 % bleach, followed by 70 % ethanol, and a propane blowtorch at the site. Bone samples several cm long were obtained using a diamond-coated dremel saw or utilising natural fractures in the bone. For the fully-aseptically-collected bone samples, the bone was sampled without first removing the surrounding matrix, although fractures in the mudstone did appear during sampling so that the samples cannot be said to have been unexposed to the air, especially prevalent in the small rib sample sent to Princeton University for analysis. Also sampled were the aseptically-excavated, but completely-exposed portions of the subterranean bone immediately next to the fully-aseptically-collected bone region (i.e., the regions of the bone fully-exposed using aseptic techniques after initial discovery of the bone in order to determine size and orientation and referred to here as aseptically-exposed). All samples were collected in autoclaved foil without applying consolidants, placed in an ice cooler kept in the shade, and brought back to the field camp freezer that evening. Surface-eroded bone from BB180 and on the same ridge above BB180, mudstone excavated from the overburden-removed area of BB180 and several cm below the weathered surface of the same ridge above BB180, and topsoil on the same ridge above BB180 were similarly aseptically-acquired and stored (i.e., sterile tools, foil, and personal wear; kept cool). In total, eight bone samples, eight sediment samples, and two soil samples were collected. Samples were transported to the Royal Tyrrell Museum of Palaeontology in a cooler. Following accession at the museum, replicates of the samples were transported on ice as logistically possible to the University of Bristol and Princeton University and stored at 4 or -80 °C, respectively, as required for analysis upon arrival (i.e., both Princeton and Bristol received a sample of fully-aseptically-collected bone, BB180 mudstone, topsoil, etc.). Samples were mailed to Princeton on ice, while samples were transported via plane to Bristol without refrigeration (maximum time unrefrigerated under 24 hr).

These aseptically-collected Dinosaur Provincial Park fossil bone, mudstone, and soil samples were compared to younger fossils and modern bone (supplemental material). In particular, chicken (*Gallus gallus domesticus*) bone was obtained frozen from a Sainsbury’s grocery store in Bristol, UK and was kept refrigerated (4 °C). This chicken bone can be safely assumed to contain high amounts of collagen. Other controls included amino acid composition data from a reference bone (fresh, modern sheep long bone) and radiocarbon data from an 82–71 Ka radiocarbon-dead bovine right femur used as a standard from the literature (Cook *et al.* 2012). Black, fossil sand tiger shark teeth (*Carcharias taurus*) eroded from Pleistocene-Holocene sediments were non-aseptically-collected from the surface of the sand on a beach in Ponte Vedra Beach, Florida, USA with no applied consolidants and were stored at room temperature. It should be noted that Florida experiences a high temperature climate relative to many samples typically studied for palaeoproteomics. Teeth samples represent a mix of dentine and enamel as opposed to normal bone tissue, with relative concentrations depending on how easily the different tissues fragmented during powdering. Technical grade humic acid was also purchased from Sigma Aldrich as a further control.

### ATR FTIR

ATR FTIR was carried out at the University of Bristol. Samples were powdered in a sterile mortar and pestle (70 % ethanol rinsed) and then demineralised in 10 mL of 0.5 M hydrochloric acid (HCl) for 5 days, with the acid replaced three times during that period by spinning in a centrifuge and pipetting off the old acid and replacing with fresh acid. After demineralisation and pipetting out the last acid volume, the samples were rinsed with 10 mL of milli-Q water and spun in a centrifuge three times, replacing the water each time. After pipetting out the last water volume, samples were freeze dried overnight.

The demineralisation products were analysed using a Nicolet iN10 MX FTIR spectrometer with a KBr beamsplitter and MCT/A detector. The sample was placed on a KBr disc, and background spectra were collected before analysing each sample for subtraction purposes. The sample contacted with a Ge tip microATR attachment that was cleaned with ethanol. Next, 128 scans were collected over a wavelength range from 675–4000 cm^-1^ at 8 cm^-1^ resolution. The spectrometer aperture windows were set to 50 µm giving an effective sample collection area of about 17 µm.

### Light microscopy, VPSEM, and EDS

The same demineralised samples that underwent ATR FTIR were subsequently analysed by VPSEM and EDS performed at the University of Bristol. Specimens were mounted onto carbon tape on standard SEM pin-stubs and were not electrically coated. A Zeiss SIGMA-HD VPSEM instrument was used in this work, with the instrument’s chamber filled with a recirculated nitrogen supply to negate against the electrical surface charge accumulation on the sample. Typical vacuum levels during analysis varied between 0.1 and 0.25 mbar. Control of the SEM, with a specified spatial resolution of 1.2 nm under such low-vacuum conditions, was performed using the microscope’s standard SmartSEM user interface. For standard sample imaging, a beam current of 1.7 nA (30 µm aperture), 15 kV accelerating voltage, and a 10 mm working height (horizontal sample; no tilt) were used. During the accompanying EDS compositional analysis of regions of interest within the sample, both the beam current and accelerating voltage were increased to 2.9 nA and 20 kV, respectively, with the sample position in the instrument remaining unchanged. An EDAX Ltd. (Amatek) Octane Plus Si-drift detector was used for the EDS analysis, with control performed through the accompanying TEAM analytical software. Collection periods of 100 s were used, with the electrically (Peltier) cooled detector operating with a dead-time of 20 % to permit for individual peak discrimination from the 30,000–40,000 counts per second incident onto the device. Elemental quantification of the spectra obtained was performed using the eZAF deconvolution and peak-fitting algorithm based upon the ratios of the differing K, L, and M X-ray emissions.

After VPSEM and EDS analysis, the same samples were imaged using light microscopy utilising a Leica DFC425 C digital camera under magnification from a Leica M205 C stereomicroscope.

### Py-GC-MS

Py-GC-MS was conducted at the School of Chemistry, University of Bristol. Sample fragments were rinsed in 70 % ethanol prior to powdering with a sterile mortar and pestle (70 % ethanol rinsed) in an attempt to remove exterior contamination. A quartz tube was loaded with ∼1 mg of the sample powder and capped with glass wool. A pyrolysis unit (Chemical Data Systems (CDS) 5200 series pyroprobe) was coupled to a gas chromatograph (GC; Agilent 6890A; Varian CPSil-5CB fused column: 0.32 mm inner diameter, 0.45 µm film thickness, 50 m length, 100 % dimethylpolysiloxane) and a double focussing mass spectrometer (ThermoElectron MAT95, ThermoElectron, Bremen; electron ionisation mode: 310 °C GC interface, 200 °C source temperature) with a 2 mL min^-1^ helium carrier gas. Samples were pyrolysed in the quartz tube (20 s, 610 °C), transferred to the GC (310 °C pyrolysis transfer line), and injected onto the GC (310 °C injector port temperature was maintained, 10:1 split ratio). The oven was programmed to heat from 50 °C (held for 4 min) to 300 °C (held for 15 min) by 4 °C min^-1^. A *m/z* range of 50–650 was scanned (one scan per second). There was a 7 min delay whereby the filament was switched off for protection against any pressure increases at the start of the run. MAT95InstCtrl (v1.3.2) was used to collect data. QualBrowser (v1.3, ThermoFinnigan, Bremen) was used to view data. Compounds were identified with the aid of the National Institute of Standards and Technology (NIST) database.

### HPLC amino acid analysis

Reversed-phase high performance liquid chromatography (RP-HPLC) for analysing amino acids was done at the University of York on samples originally sent to the University of Bristol. Samples from Dinosaur Provincial Park were transported to York on ice. Replicate sample fragments were either ethanol (70 %) rinsed prior to powdering or powdered without a rinse with a sterile mortar and pestle (also 70 % ethanol rinsed). Several mg of powder were accurately weighed into sterile 2 mL glass vials (Wheaton). Then, 7 M HCl (Aristar, analytical grade) was added, and the vials were flushed with N_2_. Hydrolysis (18 hr, 110 °C) was performed, and samples were rehydrated with a solution containing HCl (0.01 mM) and L-*homo*-arginine (LhArg) internal standard. Chiral amino acid pairs were analysed using an RP-HPLC (Agilent 1100 series; HyperSil C18 BDS column: 250 mm length, 5 µm particle size, 3 mm diameter) and fluorescence detector, using a modified method outlined by Kaufman and Manley (1998). Column temperature was controlled at 25 °C and a tertiary solvent system containing methanol, acetonitrile, and sodium acetate buffer (23 mM sodium acetate trihydrate, 1.3 µM ethylenediaminetetraacetic acid (EDTA), 1.5 mM sodium azide, adjusted to pH 6.00 ±0.01 using 10 % acetic acid and sodium hydroxide) was used. Some replicates of the samples underwent dilution to reduce peak suppression caused by high mineral content. This involved salt removal by adding 60 µL of 1M HCl to ∼2 mg of powdered sample in a 0.5 mL Eppendorf tube, sonicating for 2 mins to dissolve the powder, adding 80 µL of 1M KOH to produce a gel suspension, centrifuging for 5 mins, separating a clear solution from the gel, drying the clear supernatant by centrifugal evaporation, and finally, rehydration in 20 µL LhArg (Marc Dickinson, unpublished methods).

Principal component analysis of amino acid concentration data was run on R using the prcomp() command (scale set to ‘TRUE’ in order to normalise the data).

### Radiocarbon AMS

Radiocarbon analyses were performed at the BRAMS facility at the University of Bristol. Fossil bone samples were surface cleaned using an autoclaved razorblade to scrape their exterior surface. All samples were powdered by mortar and pestle cleaned through autoclaving and rinsing with 70 % ethanol. Samples were transferred into pre-combusted (450 °C, 5 hr) culture tubes and 10 mL of 0.5 M HCl were added to eliminate any carbonates. The HCl solution was replaced as necessary until CO2 effervescence ceased. Samples were rinsed with three washes of 10 mL MilliQ ultrapure water before freeze-drying. Samples were weighed into aluminium capsules to obtain ∼1 mg C before being combusted in an Elementar Microcube elemental analyser (also obtaining the % C by mass of the demineralised samples) and graphitised using an IonPlus AGE3 graphitisation system. The resulting graphite samples were pressed into Al cathodes and analysed using a MICADAS accelerator mass spectrometer (Laboratory of Ion Beam Physics, ETH, Zurich). All samples were blank subtracted using a bone sample known to be radiocarbon ‘dead’, the Yarnton sample from Cook *at al.* (2012).

### DNA extraction, 16S rRNA amplicon sequencing, and epifluorescence microscopy

DNA extraction and quantification (Qubit fluorometer), epifluoresence microscopy (SYTO 9/propidium iodide (PI) dual staining), and 16S rRNA amplicon sequencing were conducted at Princeton University. The bone and adjacent mudstone were processed inside a laminar flow hood after UV treatment (30 min). Specifically, the bone fragments were carefully picked out and surfaces of the fossil bones were scraped off with an autoclaved razor blade. The bone, the scrapings, and mudstone were powdered separately, with a sterile mortar and pestle after autoclaving and UV treatment. The powder from fossil bone samples were either demineralised in 0.5 M EDTA (pH = 8) or not demineralised. The EDTA demineralised bone was stained with SYTO 9 dye and propidium iodide (LIVE/DEAD BacLight Bacterial Viability Kit, Molecular Probes, USA) in the dark for 15 min. These are fluorescent dyes that intercalate between the base pairs of DNA. The stained samples were analysed using a fluorescence microscope (Olympus BX60, Japan). Live cells are stained as green whereas membrane-compromised cells fluoresce red.

Powder (0.25 g) was used to extract DNA from the bone, the scrapings and mudstone by using Power Viral Environmental RNA/DNA Isolation kit (MO BIO Laboratories, Carlsbad, CA, USA). However, the DNA yield from the mudstone was below detection (0.5 ng/mL). Therefore, a further attempt was made to extract DNA from a large amount of powder (5 g bone and 10 g mudstone) using DNeasy PowerMax Soil Kit (QIAGEN, Germanry) according to the manufacturer’s instruction. Additionally, the slurry from the EDTA demineralised bone was subjected to DNA extraction using the same large scale kit. Extracted DNA was then quantified by dsDNA HS Assay kit (Life Technologies, Carlsbad, USA) and the fluorescence was measured using a Qubit fluorometer (Invitrogen, Carlsbad, USA).

To prepare the library for 16S rRNA amplicon sequencing, DNA was used as PCR template to amplify the 16S rRNA gene V4 region using bacterial/archaeal primer 515F/806R (Caporaso *et al.* 2012). The PCR reaction condition was as follows: initial denaturation at 94 °C for 3 min; 25 or 30 cycles of denaturation at 94 °C for 45 s, annealing at 50 °C for 1 min, extension at 72 °C for 90 s and a final extension at 72 °C for 10 min. PCR product (5 µL) was loaded onto a gel to confirm the amplification by running agarose gel electrophoresis. The amplicon products were pooled to make the library and sequenced for a 150-bp paired-end reads on Illumina Hiseq 2500 housed in the Genomics Core Facility at Princeton University. The raw sequences were quality-filtered with a minimum Phred score of 30 and analysed by QIIME (Quantitative Insights Into Microbial Ecology) software package (Caporaso *et al.* 2010). Data will be uploaded onto the Sequence Read Archive (SRA) of NCBI upon publication.

## Results

### Light microscopy, VPSEM, and EDS

VPSEM and EDS of HCl demineralised, freeze-dried dinosaur bones revealed that vessels (and rare fibrous fragments) (Fig. 1A, D – E, H–J) were white, Si-dominated with O present, contained holes, and were sometimes infilled with a slightly more prominent C peak internally. Vessels occured alongside white quartz crystals, which had strong Si peaks and overall were elementally similar to the vessels, and smaller reddish minerals, originally presumed to be iron oxide or pyrite, but which had high Si content (Ba was also present).

**Figure 1.**
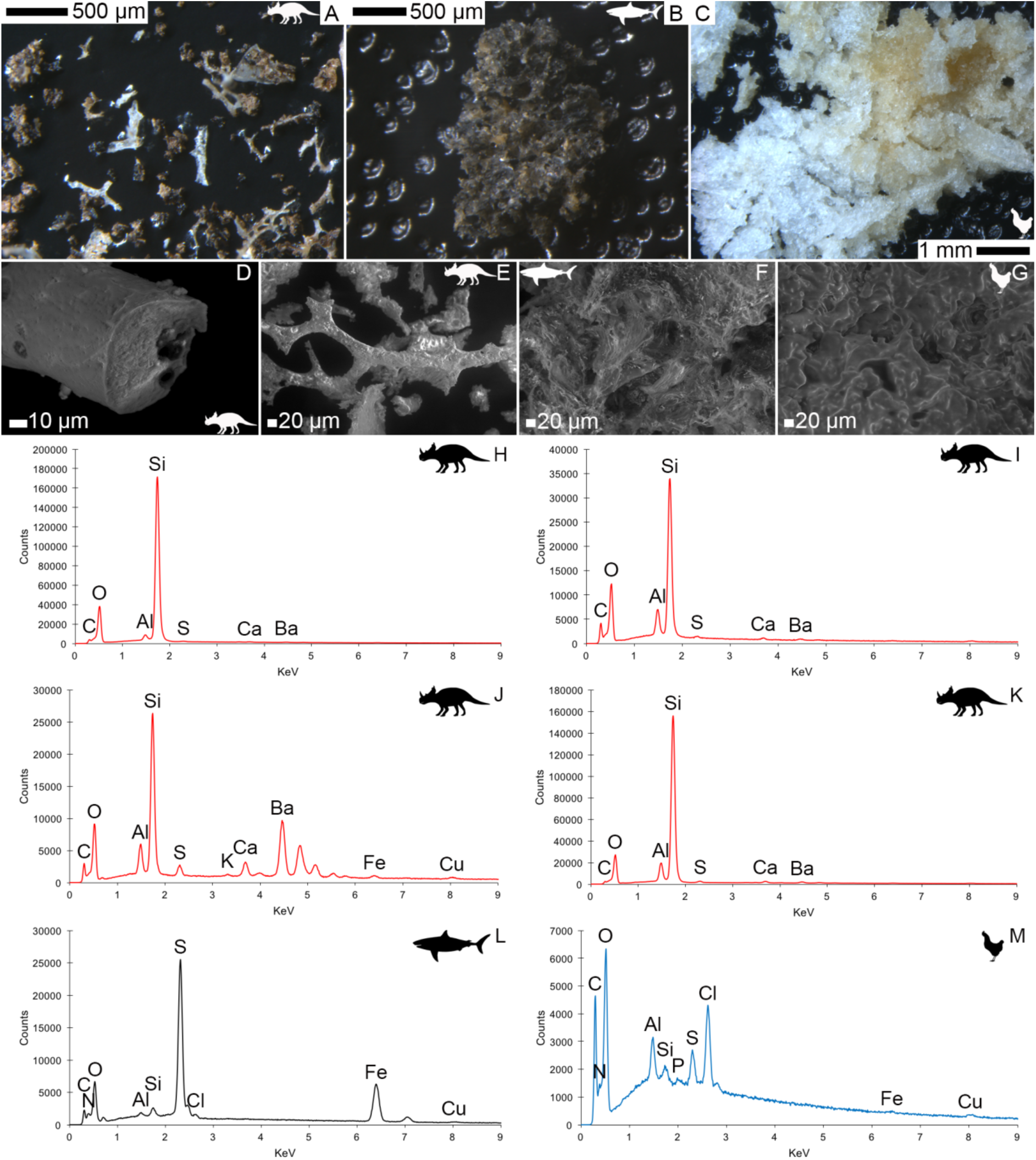
Light microscopy (A–C) and VPSEM (D–G) images and EDS spectra (H–M) of demineralised, freeze-dried samples. A–C, samples rested on carbon tape upon SEM stubs and the pitting was a result of prior VPSEM and EDS analysis. A, Centrosaurus vessels and associated minerals. B, F, L, Carcharias tooth. C, G, M, Gallus. D, infilled *Centrosaurus* vessel. E, *Centrosaurus* vessel, fibrous material along the centre of the vessel, and associated reddish minerals around the vessel. H, *Centrosaurus* vessel exterior from D. I, *Centrosaurus* vessel infilling from D. J, associated reddish mineral in *Centrosaurus*. K, *Centrosaurus* fibrous material from E. *Centrosaurus* samples are fully-aseptically-collected subterranean bone.

Demineralisation products differed from those of chicken bone (Fig. 1C, G, M) and Pleistocene-Holocene shark tooth (Fig. 1B, F, L), which were much more homogenous and consisted of large fibrous masses. These more recent samples were enriched in C, O, N, and S, but the shark tooth also had a strong Fe signature and a relatively more prominent S peak than the chicken bone. The chicken demineralisation product was white, while that of the shark tooth was black.

### ATR FTIR

ATR FTIR of a HCl demineralised, freeze-dried vessel from subterranean *Centrosaurus* bone revealed somewhat poorly-resolved, broad organic peaks (Fig. 2C) that were close in position to peaks that might be expected from various CH, CO, and amide bonds, as well as water, phosphate, and potentially carbonate and silicate bonds (Lee *et al.* 2017; also see publicly available NIST libraries). Pleistocene-Holocene shark tooth (Fig. 2B) and modern chicken bone (Fig. 2A) demineralisation products similarly revealed peaks consistent with organic and phosphatic peaks, and the chicken bone had particularly strong organic peaks relative to phosphate. Maintaining close contact of the sample to the Ge crystal was difficult, resulting in the poorly resolved peaks, especially in the shark tooth sample.

**Figure 2.**
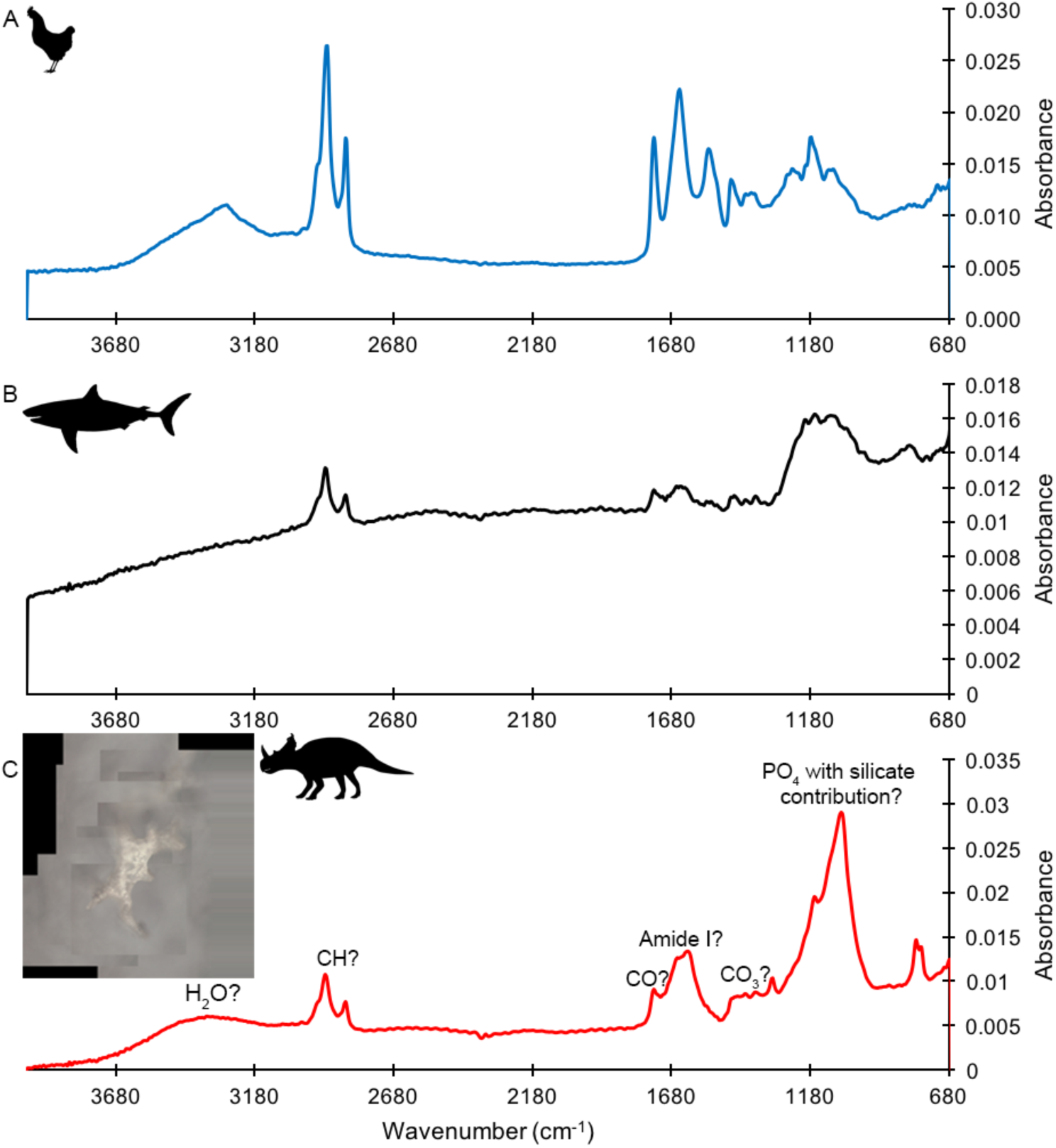
ATR FTIR spectra of demineralised, freeze-dried samples. A, Gallus. B, Carcharias tooth. C, fully-aseptically-collected subterranean *Centrosaurus* bone vessel with inset showing a composite image of the vessel that was analysed.

### Py-GC-MS

*Centrosaurus* bone had low pyrolysate concentration (Fig. 3B) as evidenced by the significant column bleed at the end of the run and contained mostly early-eluting pyrolysates. In comparison, humic acid also contained many early-eluting pyrolysates (Fig. 3D). The pyrogram for *Centrosaurus* bone does not match that of modern collagen-containing bone (Fig. 3A) and was most similar to mudstone matrix (Fig. 3C).

**Figure 3.**
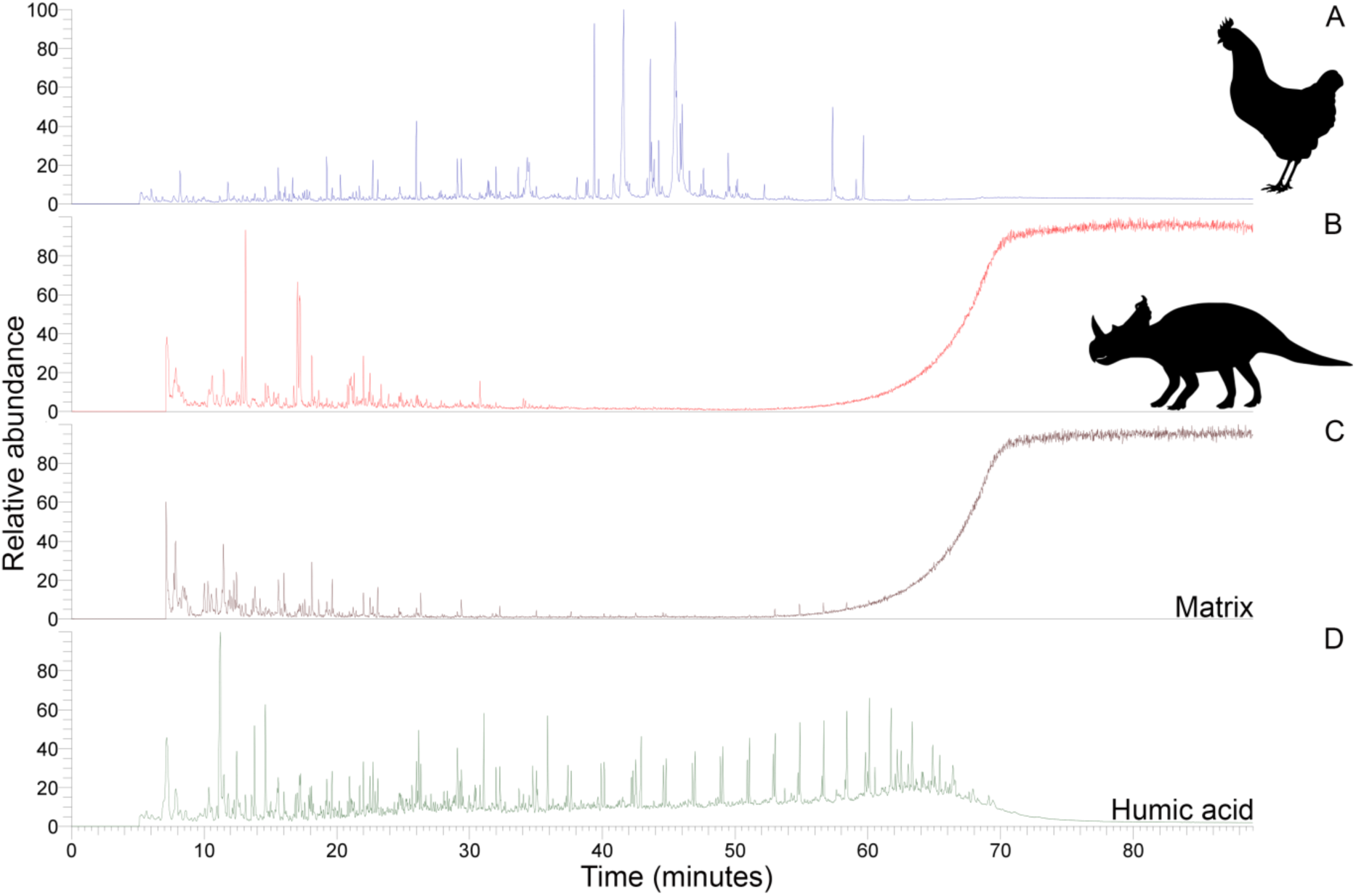
Py-GC-MS total ion chromatograms of samples ethanol rinsed before powdering. A, Gallus bone. B, fully-aseptically-collected subterranean *Centrosaurus* bone. C, adjacent mudstone matrix of subterranean *Centrosaurus* bone in B. D, humic acid (technical grade) powder.

Prominent subterranean *Centrosaurus* bone pyrolysates were indanes/indenes, alkyl benzenes, and some polycyclic aromatic hydrocarbons. Weak alkane/alkene doublets were detected in the Late Cretaceous bones (Fig. 4A–D; supplemental material). Variation in the conspicuousness of these doublets between the fully-aseptically-collected and aseptically-exposed subterranean *Centrosaurus* bone samples was apparent (Fig. 5A–D).

**Figure 4.**
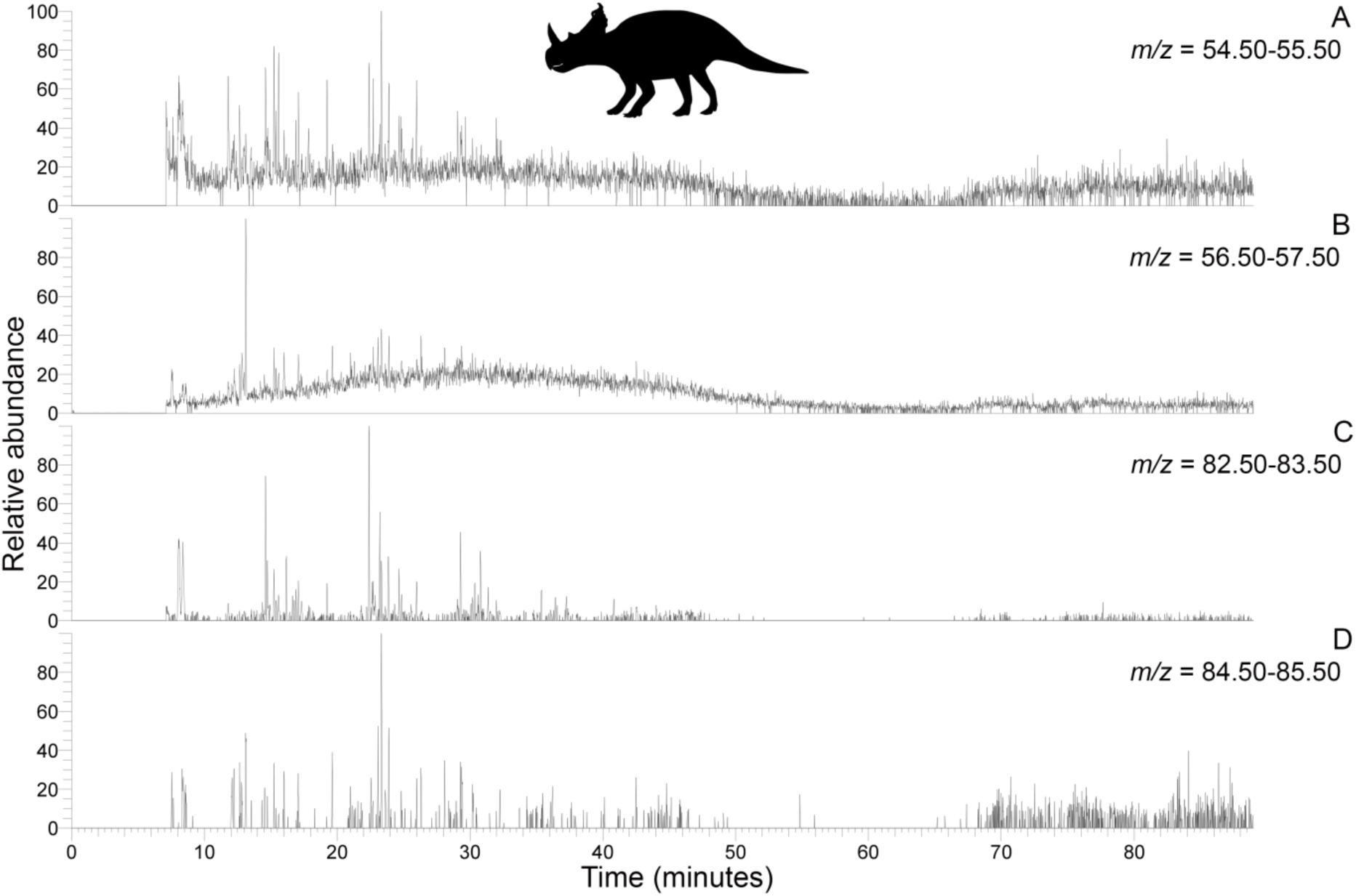
(Above). Py-GC-MS chromatograms searching for ion m/z ranges typical of alkanes and alkenes from kerogen in the fully-aseptically-collected subterranean *Centrosaurus* bone ethanol rinsed before powdering. Doublets are weakly apparent at best. A, m/z = 55. B, m/z =57. C, m/z =83. D, m/z =85.

**Figure 5.**
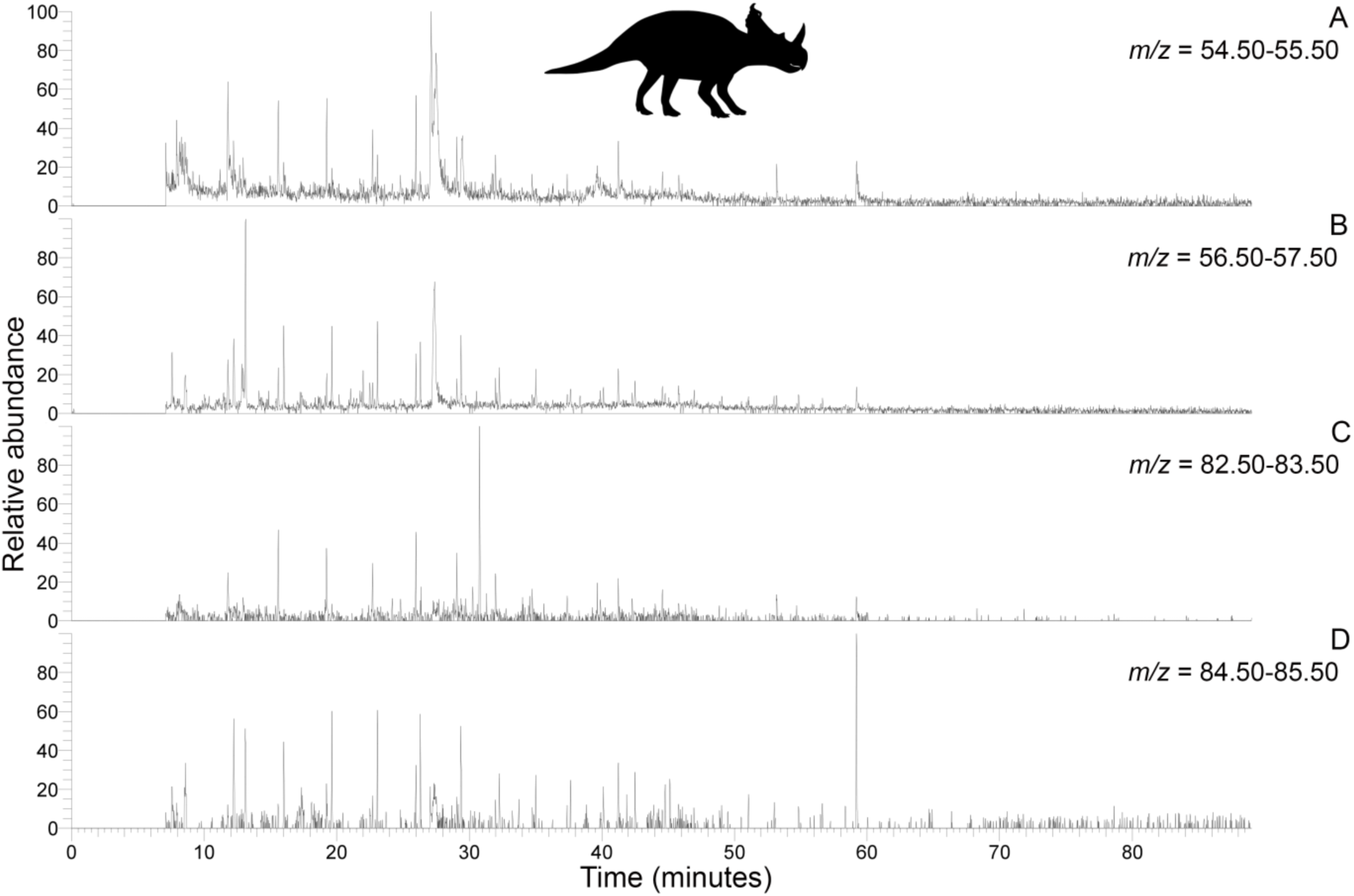
Py-GC-MS chromatograms searching for ion m/z ranges typical of alkanes and alkenes from kerogen in the aseptically-exposed subterranean *Centrosaurus* bone ethanol rinsed before powdering. Doublets are more strongly apparent than in Fig. 4. A, m/z = 55. B, m/z =57. C, m/z =83. D, m/z =85.

### HPLC amino acid analysis

Fully-aseptically-collected subterranean *Centrosaurus* bone had a total hydrolysable amino acid (THAA) compositional profile that did not match collagen (Fig. 6A, F). The fully-aseptically-collected subterranean *Centrosaurus* bone appeared to be heavily dominated by Gly. Surface-eroded Late Cretaceous bone from the same outcrop showed a different THAA compositional profile to the fully-aseptically-collected subterranean *Centrosaurus* bone, even when examining bone eroded out of the BB180 quarry itself (Fig. 6B, F). Even more interestingly, the more thoroughly treated replicates of the aseptically-exposed subterranean *Centrosaurus* bone did not match the fully-aseptically-collected subterranean bone and was similar to the surface-eroded Late Cretaceous bone in THAA compositional profile. Relative Gly concentration in surface-eroded Late Cretaceous bone was not as high as in the fully-aseptically-collected subterranean *Centrosaurus* bone, where Gly heavily dominated the compositional profile. The surface-eroded Late Cretaceous bone showed somewhat more similarity to collagen in THAA compositional profile than did the fully-aseptically-collected subterranean *Centrosaurus* bone, but ultimately did not align (Fig. 6C, F).

**Figure 6.**
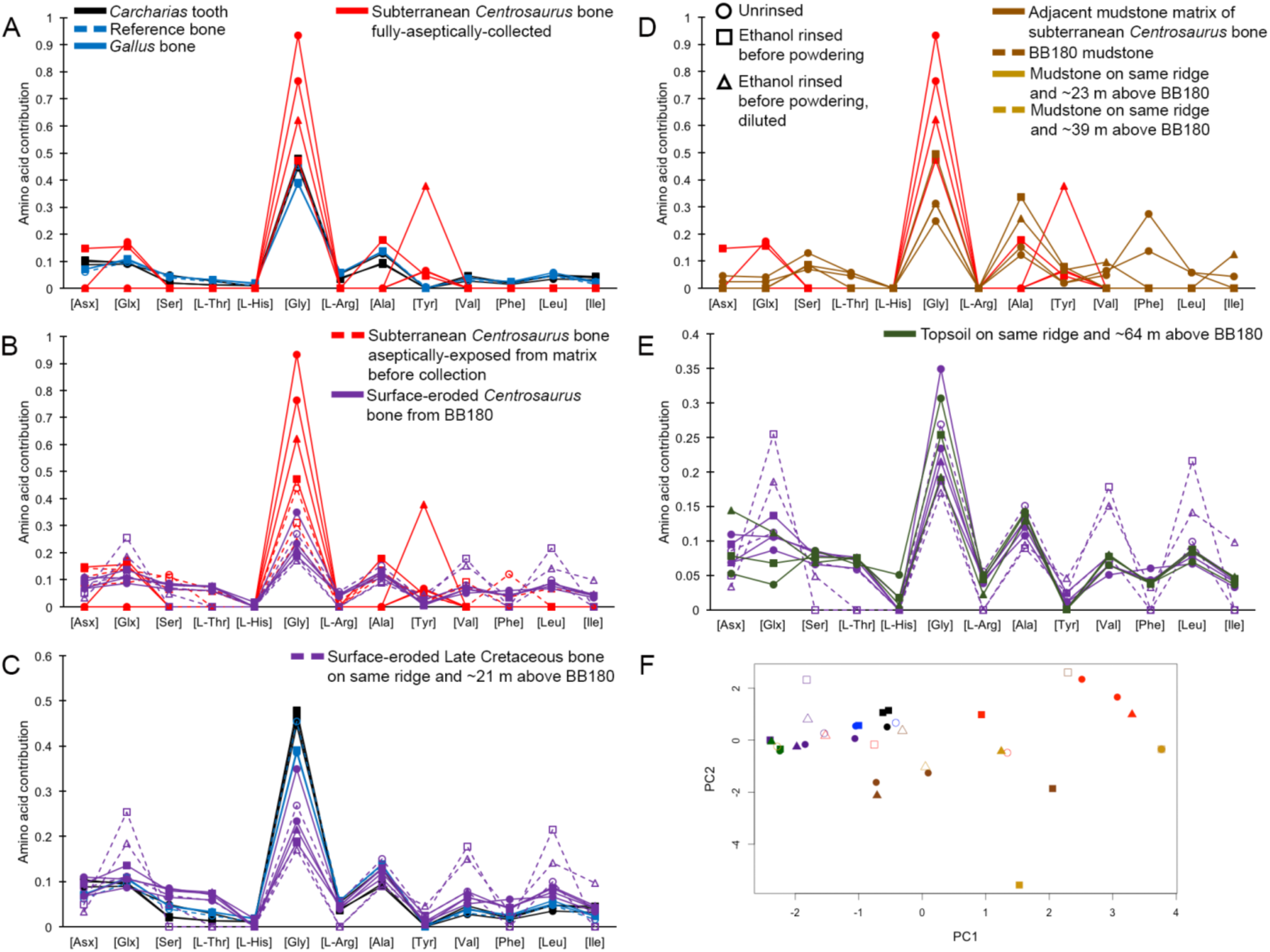
THAA compositional profiles of the samples based on amino acid percentages. A, Late Cretaceous subterranean bone (red) compared to non-aseptically-collected Pleistocene-Holocene teeth (black) and modern bone (blue). B, Late Cretaceous subterranean bone (red) compared to surface-eroded Late Cretaceous bone from the same outcrop (purple). C, surface-eroded Late Cretaceous bone (purple) compared to Pleistocene-Holocene teeth (black) and modern bone (blue). D, Late Cretaceous subterranean bone aseptically collected (red) compared to the adjacent mudstone matrix (brown). E, surface-eroded Late Cretaceous bone (purple) compared to topsoil at higher elevation (i.e., prairie level) on the same ridge (green). F, PCA on normalised amino acid percentages (see A–E legends). See supplemental material (Fig. S.21; Table S.9) for PCA summary. Colour and symbol coding is constant throughout.

Subterranean *Centrosaurus* bone had far lower THAA concentration (summed total of all amino acids measured) than did modern chicken bone (Fig. 7A), and the aseptically-exposed subterranean *Centrosaurus* bone showed higher THAA concentration than the fully-aseptically-collected subterranean *Centrosaurus* bone (Fig. 7B). Surface-eroded Late Cretaceous bone showed high variability in THAA concentrations, and at least one sample seemed to show higher THAA concentrations than subterranean *Centrosaurus* bone.

**Figure 7.**
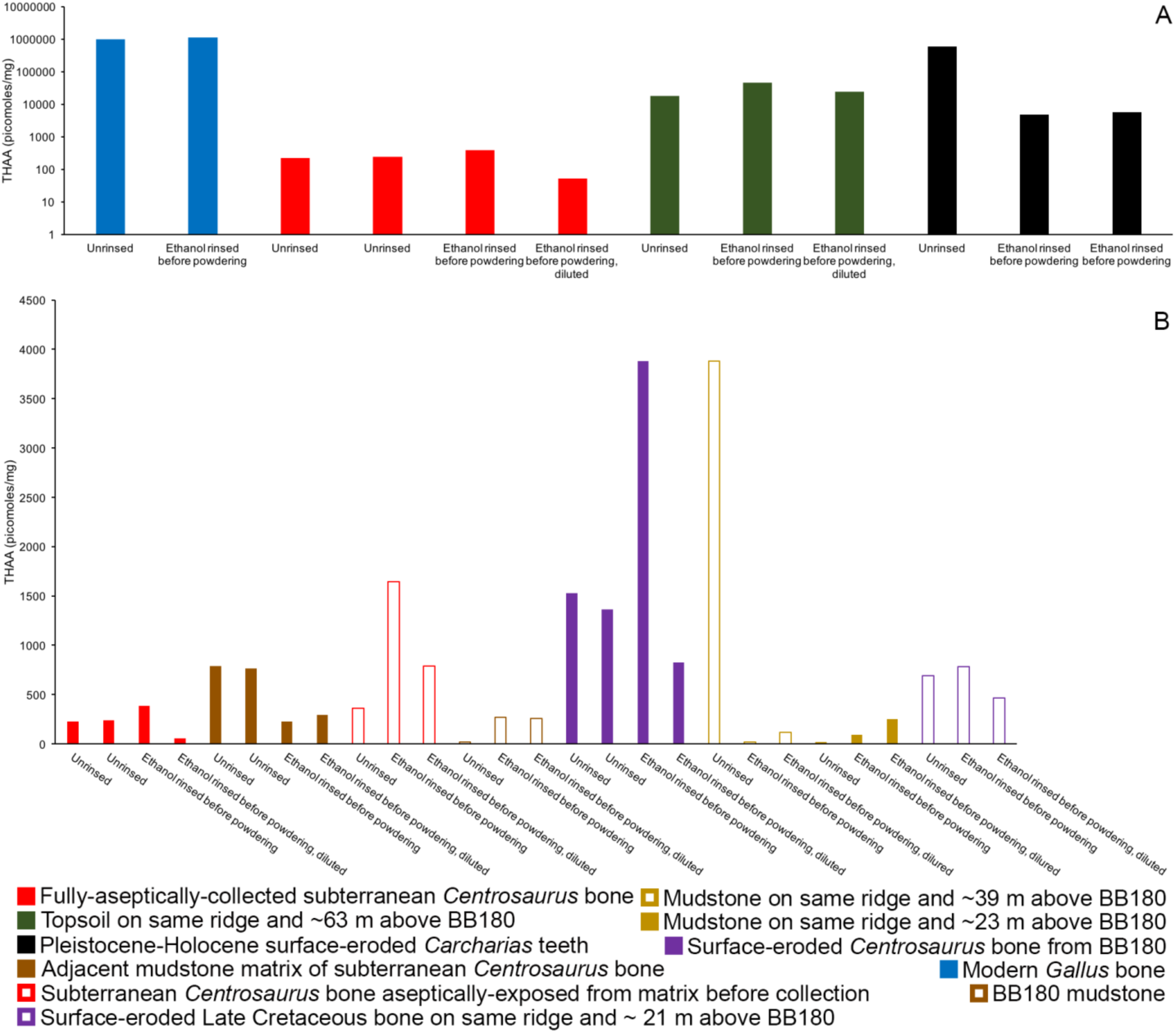
THAA concentrations (summed total of all amino acids measured) of the samples. A, logarithmic scale comparison of modern Gallus bone (blue), fully-aseptically-collected subterranean *Centrosaurus* bone (red), Pleistocene-Holocene surface-eroded shark teeth (black, with a repeated measurement for the ethanol rinsed sample), and topsoil on same ridge and ∼64 m above BB180 (green). B, comparison between fossil Late Cretaceous bone and mudstone. Fully-aseptically-collected subterranean *Centrosaurus* bone (solid red), adjacent mudstone matrix of subterranean *Centrosaurus* bone (solid brown), aseptically-exposed subterranean *Centrosaurus* bone (open red), BB180 mudstone (open brown), surface-eroded *Centrosaurus* bone from BB180 (solid purple), mudstone on same ridge and ∼39 m above BB180 (open tan), mudstone on same ridge and ∼23 m above BB180 (solid tan), and surface-eroded Late Cretaceous bone on same ridge and ∼21 m above BB180 (open purple). Diluted replicates likely provide the most accurate measurements given the peak reduction present in the non-diluted replicates.

Late Cretaceous bone was L-amino acid dominated when such amino acids were above detection limit (Table 1). Surface-eroded Late Cretaceous fossil bone seemed to show slightly more variability in D/L values than the subterranean bone samples. Similar to the samples described here, other, non-aseptically-collected, room-temperature-stored Jurassic and Cretaceous surface-eroded bones have low amino acid concentrations and lack significant concentrations of D-amino acids (supplemental material).

**Table 1.**
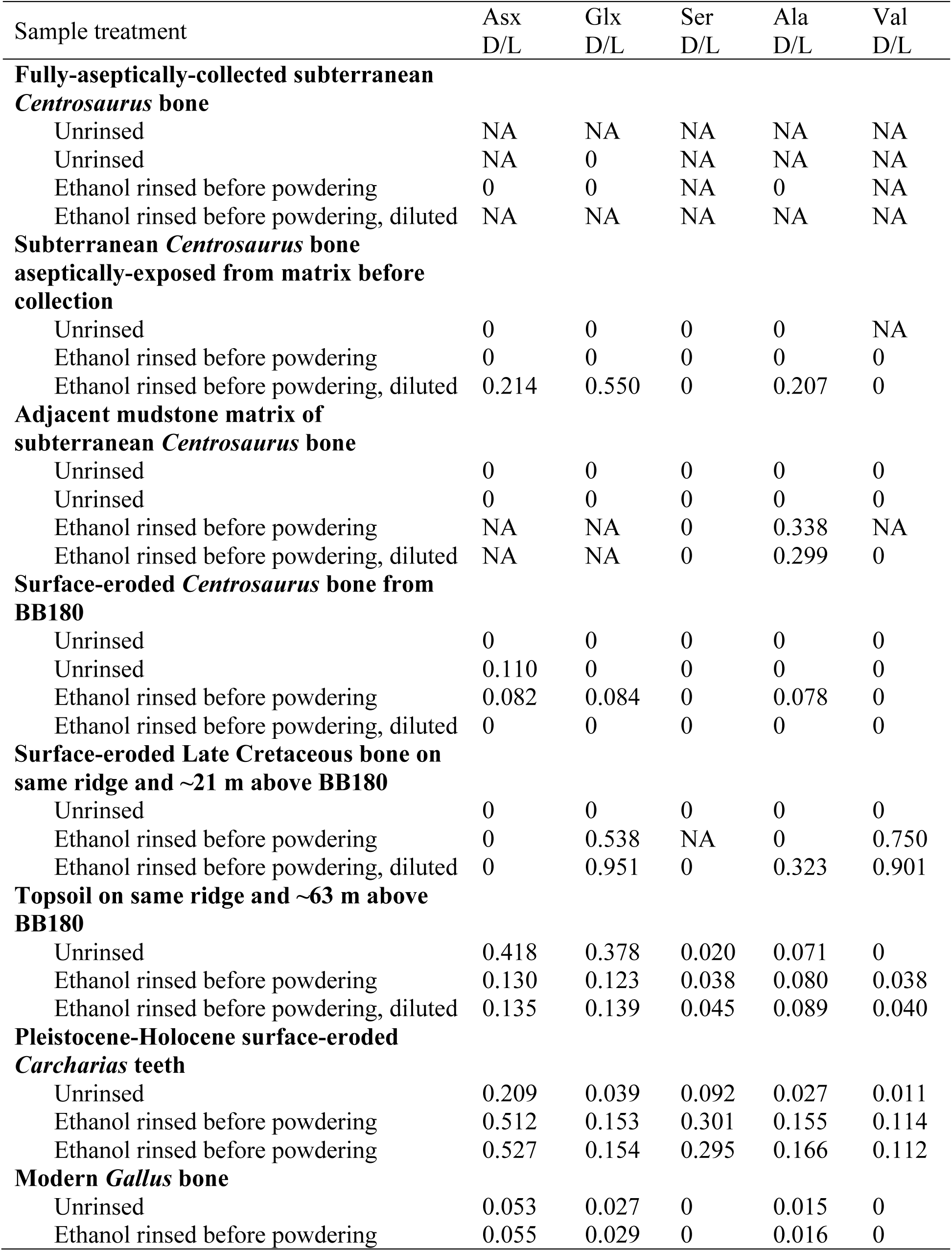
Comparison of Late Cretaceous, Pleistocene-Holocene, and modern amino acid racemisation values. NA indicates that amino acid concentration was below detection limit.

The adjacent mudstone matrix did not match the subterranean *Centrosaurus* bone in THAA compositional profile (Fig. 6D, F). Some of the surface-eroded Late Cretaceous bone showed similarity to topsoil at the prairie level above the outcrop in THAA compositional profile (Fig. 6E, F). Although mudstone and fully-aseptically-collected subterranean *Centrosaurus* bone did not tend to overlap in THAA compositional profile, they differed from collagen in diametrically different ways than did the topsoil and surface-eroded Late Cretaceous bone (Fig. 6F). The fully-aseptically-collected subterranean *Centrosaurus* bone tended to have higher Gly and Tyr and lower Glx, Leu, Asx, Val, Ile, and L-Thr than collagen. Mudstone tended to have higher Tyr, Ala, and Ser than collagen. Topsoil and surface-eroded Late Cretaceous bones (as well as the more thoroughly treated replicates of the aseptically-exposed subterranean *Centrosaurus* bone) tended to have higher Glx, Leu, Asx, Val, Ile, and L-Thr and lower Gly and Tyr than collagen. The greatest variation between the samples of this study was in relative Gly and Tyr concentrations, although Tyr variation was heavily skewed by two samples (one mudstone and one fully-aseptically-collected subterranean *Centrosaurus* bone) with unusually high concentrations. Most of the consistently observed variation occured in Gly relative concentrations, and this metric appeared somewhat able to discriminate between 1) fully-aseptically-collected subterranean *Centrosaurus* bone, 2) modern bone and Pleistocene-Holocene surface-eroded shark teeth, and 3) topsoil, mudstone, and surface-eroded Late Cretaceous bone (supplemental material).

Topsoil showed greater THAA concentration than subterranean and surface-eroded *Centrosaurus* bones, but not as high as modern chicken bones (Fig. 7A). Mudstone tended to have very low THAA concentration, even compared to some of the Late Cretaceous bone samples (Fig. 7B); however, one unrinsed replicate of mudstone ∼39 m above BB180 was an outlier, potentially representing contamination or collection of pure mudstone alongside some larger organic detritus. The highest THAA concentrations in mudstone tended to be observed in the adjacent mudstone matrix to the subterranean *Centrosaurus* bone. When amino acids were above detection limit, mudstone was L-amino acid dominated like the Late Cretaceous bone (Table 1). Topsoil, on the other hand, showed moderate levels of racemisation.

Pleistocene-Holocene surface-eroded shark teeth had THAA compositional profiles that strongly matched collagen (Fig. 6A, C, F) and fairly high amino acid concentration with THAA concentrations between those of subterranean *Centrosaurus* bone and modern chicken bone (Fig. 7A). Pleistocene-Holocene surface-eroded shark teeth, unlike the Late Cretaceous bone and mudstone, had very high racemisation (Table 1), even more so than the topsoil sample. Ethanol rinsing appeared to lower amino acid concentration in the shark teeth but did not affect THAA compositional profile (Figs. 6A, C, F, 7A).

### Radiocarbon AMS

Total organic carbon (TOC) content was higher in the subterranean and surface-eroded *Centrosaurus* bone than the matrix, even the directly adjacent matrix, and was comparable to that found in the topsoil (Table 2). However, the organic carbon content in the *Centrosaurus* bones was significantly lower than the 82–71 Ka Yarnton bovine bone sample known to contain well-preserved (radiocarbon-dead) collagen (Cook *et al.* 2012). TOC in the *Centrosaurus* bone was not found to be radiocarbon dead, but did exhibit lower F^14^C values than both the mudstone and especially the topsoil. Assuming all endogenous bone C is radiocarbon ‘dead’, based on these F^14^C values, a simple 2-end-member mixing model would suggest that ∼26 % of the C in subterranean *Centrosaurus* bone originates in the adjacent mudstone matrix (supplemental material).

**Table 2.**
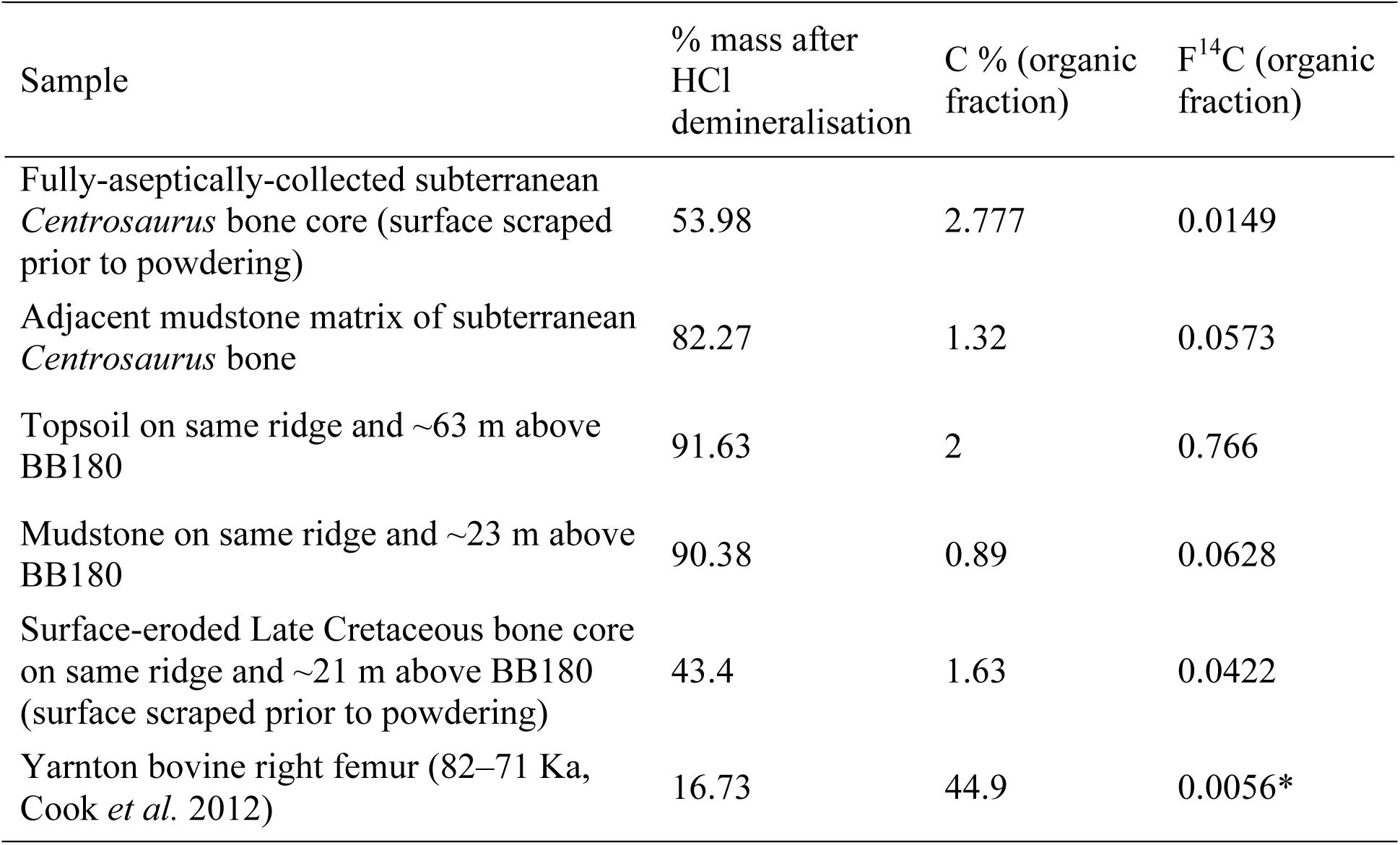
Carbon data from Late Cretaceous fossil bone, mudstone, topsoil, and younger bone. *This sample was used for blank correction in the AMS analyses, therefore this value is not blank-subtracted.

### Fluorescence microscopy, DNA extraction and 16S rRNA amplicon sequencing

DNA concentration was about 50 times higher in subterranean *Centrosaurus* bone than in adjacent mudstone matrix (Table 3; supplemental material). PI staining for DNA on EDTA demineralised *Centrosaurus* bone revealed multi-cell aggregates forming organic vessel and conglomerate structures that fluoresce red (Fig. 8A–D).

16S rRNA amplicon sequencing revealed the predominance of Actinobacteria and Proteobacteria in subterranean *Centrosaurus* bone (Fig. 9). Sequences affiliated with classes Nitriliruptoria and Deltaproteobacteria were more abundant relative to adjacent mudstone or even the surface scrapings from the bone itself. The majority of the sequences within Deltaproteobacteria were identified as belonging to the family Desulfurellaceae, which contains some sulfur-respiring species (sulfur-, rather than sulfate-reducing). However, the short reads prevented species level identification. In *Centrosaurus* bones, about 30 % of sequences were phylogenetically close to the genus *Euzebya*.

**Table 3.** DNA concentrations in mudstone matrix and bone quantified with Qubit fluorometry.

| Sample | Average DNA concentration (ng/μL) | Total DNA (ng) | DNA per 1 g of bone or mudstone (ng/g) |
| --- | --- | --- | --- |
| Fully-aseptically-collected subterranean *Centrosaurus* bone core (surface scraped prior to powdering) | 0.793 | 3965 | 793 |
| Adjacent mudstone matrix of subterranean *Centrosaurus* bone | 0.0328 | 164 | 16.4 |
| Laboratory blank | Below detection (<0.0005 ng/μL) | | |

**Figure 8.**
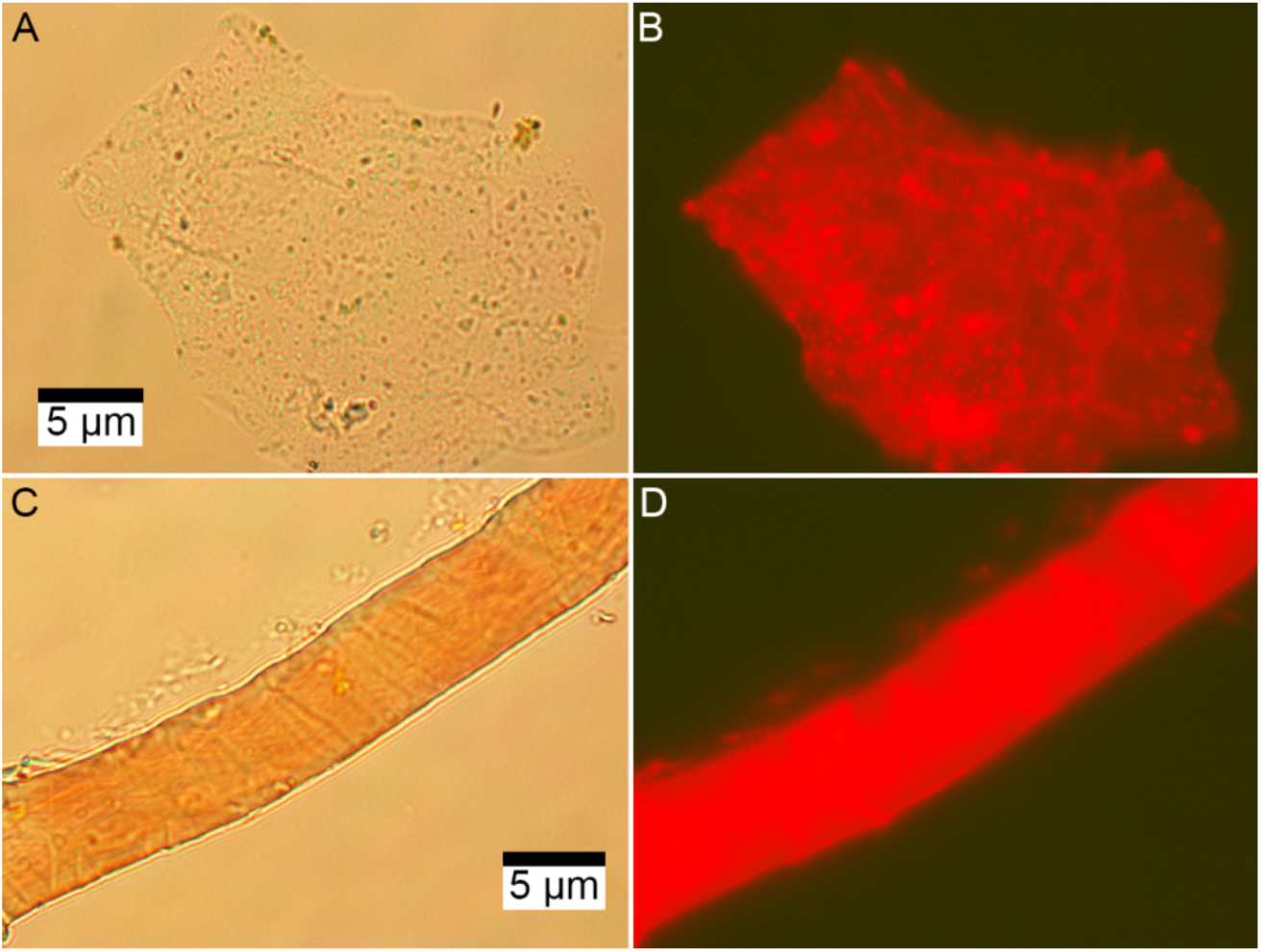
Microscopic images of EDTA demineralised, PI stained fully-aseptically-collected subterranean *Centrosaurus* bone. A–B, fibrous material. C–D, vessel. A, C, Transmission light. B, D, Fluorescence.

**Figure 9.**
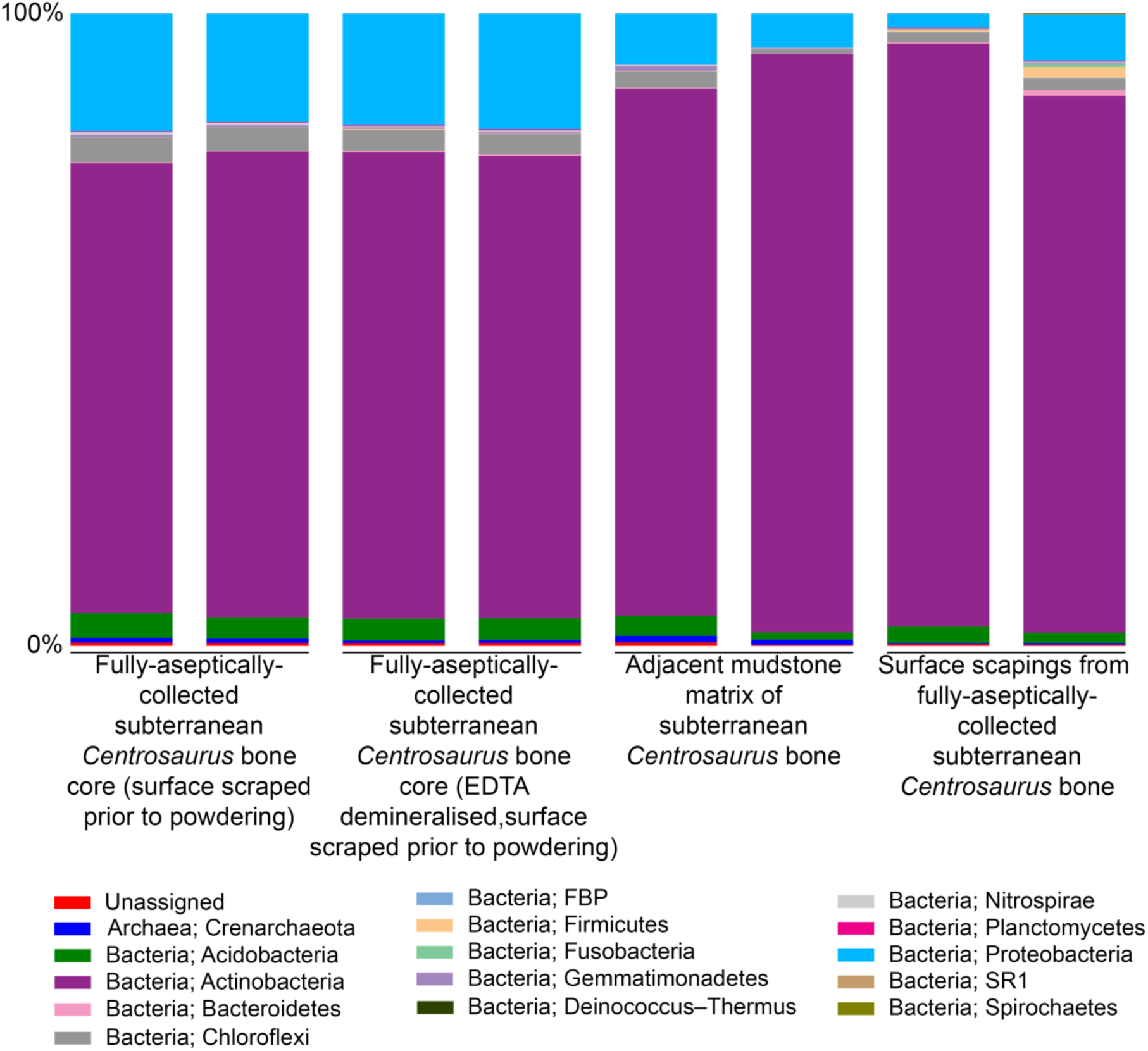
Comparison of microbial community (phylum level) from fully-aseptically-collected subterranean *Centrosaurus* bone and adjacent mudstone matrix. There are two replicates per sample.

## Discussion

### Light microscopy, VPSEM, and EDS

Demineralisation products of dinosaur fossil bone differ structurally and elementally from the Pleistocene-Holocene and modern samples when examined using light microscopy and VPSEM. Occasional infilling of the dinosaur bone vessels with greater C concentration in the interior compared to the exterior of the vessel is consistent with a growing biofilm. The Si dominance of the demineralisation products from the dinosaur bone likely suggest that they are at least partly silicified. HCl demineralisation (especially the relatively intensive demineralisation used on the samples that underwent microscopy, EDS, and ATR FTIR) may favour mineralised biofilm retrieval assuming that low pH might degrade organically-preserved biofilms, explaining why all of the observed demineralisation products of the dinosaur bone have high Si content. If this is the case then it would indicate that the original organics are significantly more susceptible to acid (i.e., of different composition) than the organic masses in the identically treated younger bone samples, which survive well. It seems likely that mineral infilling in the *Centrosaurus* bone is largely of silicates, which have partly replaced the originally organic vessel-like structures, as well as, potentially, some barite or gypsum with minimal iron oxide or pyrite. Some of these inorganic compounds might contribute to the colour of the fossils. The fibrous material may be silicified biofilm with a collagenous texture imprint from the surrounding apatite matrix or may simply be a small misinterpreted quartz crystal. A silicified biofilm might be a result of unique environmental conditions (either early or late in the taphonomic history) or microbial communities that these fossils experienced. Therefore, examining fossils from different localities, climates, lithologies, and taphonomic histories is vital to understanding variation in how biofilms in fossil bone might be mineralised.

The Pleistocene-Holocene shark tooth and modern chicken bone demineralise to reveal large organic masses (i.e., rich in C and O) consistent with collagen protein as evidenced by discernable N and S peaks, unlike the much older dinosaur bone demineralisation products. However, the high Fe content in the shark tooth suggests some taphonomic mineral accumulation (e.g., iron oxide or pyrite) and may explain some of the dark discolouration in the teeth, potentially alongside a browning effect caused by the taphonomic formation of melanoidins, advanced glycation end products, and similar condensation products (Wiemann *et al.* 2016). The relatively more pronounced S peak in the shark tooth as compared to the chicken bone might indicate sulphurisation of the collagen protein or some other taphonomic incorporation of inorganic S from the environment into the tooth, the latter being consistent with pyrite. After all, the teeth are the only fossils in this study to derive from a marine depositional environment, so the potential for pyrite formation under euxinic conditions, for example, would not be surprising. Low pressure conditions of VPSEM and EDS, as well as charging during these analyses, may have affected subsequent light microscopy observation, but this is mitigated by the fact that light microscopy was done under a comparative framework between the samples.

### ATR FTIR

The ATR FTIR results here showing various organic bonds could be taken by some to be consistent with supposed dinosaur collagen. Similar, albeit higher-resolution, peaks to those detected here are used as evidence for purported dinosaur collagen (Lee *et al.* 2017), but, it should be noted that such results are not conclusive of collagen. Detection of peaks such as those associated with amide bonds may not necessarily derive from proteins, as amide bonds are not specific to proteins and can be found in protein degradation products such as diketopiperazines (Saitta *et al.* 2017). CH and CO bonds are even more widely distributed through organic molecules. Some researchers have indeed attempted to observe how ATR FTIR spectra of bone collagen is modified when carbonaceous contamination (e.g., applied organics like consolidants, humic acids, or soil carbonate) is present (D’Elia *et al.* 2007), but it can be tempting for taphonomists to observe organics peaks in such spectra and attribute them to endogenous protein. Even if such bonds are from proteins, without deconvolving peaks to produce fingerprints of protein secondary structure (Byler & Susi 1986), one cannot say from the presence of such organic bonds alone that the protein is collagen, let alone endogenous or ancient. Despite the strong demineralisation treatment, it appears that some phosphate remained in the samples, as evidenced by the presence of peaks consistent with phosphate in the spectra. It has been shown experimentally and theoretically that variation in the phosphate bands derived from ATR FTIR of bone can be affected by bone collagen content, with low-frequency symmetry of the phosphate peaks more apparent in bone containing lower amounts of collagen (Aufort *et al.* 2018). The observation of sharper, more symmetric phosphate peaks in the *Centrosaurus* bone compared to the younger bone might suggest lower relative collagen content. However, it should be noted that the described pattern in phosphate peak alteration was observed using a diamond ATR, and this method can result in differences in spectra from those made using Ge ATR as was done here due to different refractive indices of the crystals (Aufort *et al.* 2016), so such a comparison may be inappropriate. Additionally, it would be advisable to obtain ATR FTIR data from non-demineralised samples before trying to interpret the results here. Regardless, discussion of symmetry in the phosphate peaks should be had on any future papers that attempt to use ATR FTIR data as evidence for purported Mesozoic collagen. Future work on the specimens analysed here should also attempt ATR FTIR mapping on polished sections to examine how peaks are spatially distributed, perhaps in combination with time-of-flight secondary ion mass spectrometry (TOF-SIMS).

### Py-GC-MS

The dinosaur fossil bones show greater chemical resemblance to mudstone than to fresh, modern bone and appear somewhat low in organics relative to fresh, modern bone. Although indanes and indenes have a similar carbon skeletal structure to heterocyclic indoles known to derive from protein pyrolysis, and benzenes are protein pyrolysates, these detected pyrolysates are relatively simple and are not as indicative of high proteinaceous content as would be amides, succinimides, and piperazines (Saitta *et al.* 2017). Humic acids are common in soils and contain low molecular weight, aromatic components (Hatcher *et al.* 1981; Sutton & Sposito 2005), meaning that the early-eluting, presumably more volatile (i.e., lower boiling point), peaks from the *Centrosaurus* bone may come from sources other than proteins.

Evidence of kerogen (in the form of alkane/alkene doublets) has been detected in the *Centrosaurus* bone, but the doublets are very weak. Variation in the conspicuousness of the doublets between the fully-aseptically-collected and aseptically-exposed subterranean *Centrosaurus* bone samples is likely representative of intra-bone variation in kerogen content rather than contamination since a strong kerogen signature is not likely to result from exposure to air or the sterilised excavating equipment. Future analyses should examine these samples by mass spectrometry under selected ion monitoring (SIM) scanning mode with comparison to an alkane/alkene standard or modify extraction methods prior to analysis in order to more clearly observe these doublets. Kerogen forming from *in situ* polymerisation of endogenous labile lipids such as cell membranes would not be expected to preserve the tubular shape of ‘soft tissues’ such as vessels in bone with high fidelity since initial hydrolytic cleavage from a hydrophilic group will eliminate the amphiphatic nature of these molecules and make them incapable of retaining their bilayer configuration in aqueous solution (Rand & Parsegian 1989). Thus, cell membranes would lose their structure, and presumably, the tubular structure of vessels would be lost as well. The possibility that the resulting kerogen could contribute to a non-tubular, low-resolution organic mould of such ‘soft tissues’ formed in the cavities of the bone’s inorganic matrix should be consider in cases in which bone demineralisation products are not mineralised. However, EDS revealed mineralisation of the structures studied here, common observation for such ‘soft tissue’ remains (Schweitzer *et al.* 2014), which is more consistent with a biofilm origin (Schultze-Lam *et al.* 1996; Decho 2010) rather than a kerogen origin. Furthermore, kerogen-like aliphatic alkanes and alkenes have been detected using Py-GC-MS from the humic fraction of soil, potentially derived from stable plant biopolymers like cuticle (Saiz-Jimenez & De Leeuw 1987), so kerogen in the fossil bone could be derived, at least partly, from soil contaminants rather than being derived from endogenous lipids.

### HPLC amino acid analysis

Amino acids in the dinosaur bone are recent and like derive from proteins other than collagenous. Low amino acid concentrations and THAA compositional profiles that do not match collagen, despite high Gly content, suggest that endogenous collagen protein has been lost from the dinosaur fossils. Although one might expect changes in the THAA compositional profile as a result taphonomic alteration and preferential loss of less stable amino acids, the lack of a THAA compositional profile that matches collagen in the dinosaur bone at the very least suggests that collagen is not well-preserved, and, therefore, structural preservation of bone collagen (as in the shark tooth) or the preservation of epitopes for antibodies are likely unsupported since these would require high levels of sequence and higher-order structural preservation. The dominance of L-amino acids in the dinosaur fossils suggests recent amino acid input. There appears to be a trend towards greater abundance of amino acids in the dinosaur bone compared to the mudstone, suggesting that the fossil bone might be preferentially concentrated in exogenous amino acids.

Surface-eroded and fully-aseptically-collected subterranean Late Cretaceous bone differ from collagen in opposite ways with respect to their THAA compositional profiles. This might suggest that subterranean bone provides a different microenvironment than surface bone, perhaps largely driven by differences in oxygen availability, and thereby containing a different microbial community. This is further evidenced by the fact that the surface-eroded Late Cretaceous bone THAA compositional profile more closely matches topsoil while the fully-aseptically-collected subterranean bone plots more closely to, although still distinct from, mudstone. This might suggest that surface-eroded bone supports a microbial community more similar to other surface communities, such as topsoil, while the subterranean bone contains a more unique community. High variability in THAA concentration in the surface-eroded Late Cretaceous bones is not surprising given that one of the surface-eroded fragments appeared to have relatively higher mineral infiltration, evidenced by greater difficulty in powdering the sample, suggesting the potential for different microenvironments inside the bone and different carrying capacities for a microbial community. One surface-eroded sample came from an active palaeontological quarry and was likely exposed to high levels of human-induced contamination as a result. The fact that one surface-eroded Late Cretaceous bone appeared to have higher THAA concentration than did the subterranean bone further suggests that bone can be colonised by exogenous microbes since surface exposure would be expected to result in adverse conditions for any surviving endogenous proteins. However, such comparisons should be done cautiously given the small sample size of this study, an aspect that should be kept in mind throughout the analyses performed here and in other studies. The most surprising result might be that the more thoroughly treated replicates of the aseptically-exposed subterranean *Centrosaurus* bone sample had a THAA compositional profile more closely matching surface-eroded bone than the fully-aseptically-collected subterranean bone and also had elevated THAA concentration compared to the fully-aseptically-collected subterranean bone, suggesting that even relatively brief aerial exposure might lead to rapid contamination on the subterranean microbial community by surface microbes.

The fact that the adjacent mudstone matrix of the subterranean bone tended to show greater THAA concentration than the other mudstone samples may indicate that bone provides a nutrient source that encourages microbial proliferation.

In contrast to the Late Cretaceous bone, the much younger shark teeth from the Pleistocene-Holocene have relatively high amino acid concentrations whose THAA compositional profiles match collagen, suggesting the presence of collagen protein. Since ethanol rinsing did not change the THAA compositional profile of shark teeth, this suggests that the majority of the organics are deriving from insoluble collagen with fairly well preserved higher-order structure, rather than highly fragmented peptides with greater mobility. This observation is consistent with the results from light microscopy and VPSEM. The shark teeth also have relatively high racemisation, a testament to the antiquity of the amino acids as would be expected from endogenous collagen.

### Radiocarbon AMS

The fact that the C in the dinosaur bone is not radiocarbon dead suggests an influx of more modern C (i.e., not radiocarbon dead) into the fossil. However, lower F^14^C in the dinosaur bone compared to the mudstone or topsoil suggests some biologically-inaccessible, old, and possibly endogenous C within the fossils. One possibility for this is kerogen derived from *in situ* polymerisation of endogenous dinosaur labile lipids, although such kerogen alkanes/alkenes have only been weakly detected in the *Centrosaurus* bones through Py-GC-MS, potentially due to methodology rather than low concentration. Exogenous C could also become metabolically inaccessible in bone through biofilm mineralisation, as suggested by the EDS data, allowing for ^14^C depletion. Additionally, biofilm formation and proliferation in bones could trap mobile organic C from sediment and groundwater at a rate faster than C exits the bone when not colonised by a biofilm. This would allow for a lower F^14^C steady state to be reached during the time it takes C outflux to increase in order to match C influx, assuming a simple 1-box model. Perhaps a combination of these three mechanisms influences F^14^C.

### Fluorescence microscopy, DNA extraction and 16S rRNA amplicon sequencing

Analyses of nucleic acids reveal a diverse, unique microbial community within the dinosaur bone even when compared to the immediate mudstone matrix or the exterior surface of the bone, as evidenced by a strong enrichment in DNA and distinct community composition in the bone. The microbial community from the EDTA demineralised bone was similar to that of the non-demineralised bone, a useful fact to be aware of since EDTA can be used as the demineralising agent to study the ‘soft tissues’ of fossil bone (Cleland *et al.* 2012). Thus, bone samples treated with common methods of demineralisation in other taphonomic studies (e.g., antibody-based studies) are also amenable for nucleic acid analyses that can be used to help test the endogeneity of organics (i.e., whether there are microbes present that could possibly explain the presence of specific organics).

PI staining of extra-nuclei tissues is potentially due to cell lysis and DNA release during demineralisation. Although further work is required to determine if the detected Deltaproteobacteria lineages can reduce sulfur to sulfide, these results may indicate that the microenvironment inside the fossil bone creates anaerobic niches to support anaerobic metabolism. Their abundance might also explain pyrite and iron oxide framboid observations in fossil bone, which can superficially resemble erythrocytes (Martill & Unwin 1997; Kaye *et al.* 2008).

## Conclusions

Previous studies have often assumed that fossil bones are biologically inert, closed systems and that organics within dinosaur bone are therefore likely to be endogenous. However, even prior to the sort of chemical analysis conducted here, simple field observations suggest otherwise. In Dinosaur Provincial Park and many other localities, fossil bone is frequently colonised by lichen on the surface or penetrated by plant roots in the subsurface. This forces researchers to consider that subsurface biota (e.g., plant roots, fungi, animals, protists, and bacteria) could contaminate bone. Given that fungi can produce collagen (Celerin *et al.* 1996), the need to rule out exogenous sources of organics in fossil bone is made all the greater. Even deeply buried bone might have the potential to be biologically active, given that microbes can thrive kilometres deep into the crust (Lin *et al.* 2006). The analyses presented here support the idea that far from being biologically ‘dead’, fossil bone supports a diverse, active, and specialised microbial community. Given this, it is necessary to rule out the hypothesis of subsurface contamination before concluding that fossils preserve geochemically unstable endogenous organics, like proteins.

We detected no evidence of endogenous proteins in the bone studied here and are therefore unable to replicate the claims of protein survival deep into the fossil record, such as the Mesozoic (Pawlicki *et al.* 1966; Schweitzer *et al.* 2005a, 2007a, 2007b, 2008, 2009, 2013, 2014, 2016; Asara *et al.* 2007; Organ *et al.* 2008; Schweitzer 2011; Bertazzo *et al.* 2015; Cleland *et al.* 2015; Schroeter *et al.* 2017). In contrast, recent Pleistocene-Holocene material can exhibit clear evidence along multiple lines of investigation for endogenous, ancient collagen, even when the fossil (dentine/enamel in this case) is stained black, indicating taphonomic alteration, and the sample is found exhumed in a warm climate and not treated with aseptic techniques. Detection of particular organic signatures in fossils (e.g., amide bands in FTIR) requires corroborating evidence before claims of ancient proteins can be substantiated. In addition to reliable markers of general protein presence (e.g., amide, succinimide, or piperazine pyrolysates), evidence is required to identify the type of protein (i.e., amino acid composition or sequence) as well demonstrate its endogenous origin (e.g., localisation) and age (i.e., degree of degradation as revealed by amino acid racemisation, deamidation, or peptide length/degree of hydrolysis). Degradation of collagen polypeptides follows a pattern of gradual hydrolysis of amino acids at the terminal ends followed by catastrophic degradation and rapid hydrolysis due to rupture of the triple helix quaternary structure, making the resulting gelatinous fragments susceptible to rapid leaching from the bone or microbial degradation (Collins *et al.* 1995, 2009; Dobberstein *et al.* 2009). It might therefore be suspected that if ancient collagen does indeed persist in a fossil bone, then such preservation would more often than not provide clear, strong structural and chemical signatures like that in the Pleistocene-Holocene shark teeth. Recently it has been suggested that techniques that do not provide information on the precise sequence or post-translational modification of peptides, such as Py-GC-MS or HPLC amino acid analysis, are outdated for palaeoproteomic studies (Cleland & Schroeter 2018). This might be the case when samples are very young and from cold environments, in which case, more precise mass spectrometry analyses such as liquid chromatography-tandem mass spectrometry might be employed early on in the course of research with elevated confidence that ancient proteins are capable of persisting in the sample. However, our results here suggest that techniques like Py-GC-MS or HPLC that give more general information on protein presence versus absence or general amino acid composition should be considered frontline approaches when dealing with fossils of significant age and/or thermal maturity (e.g., Demarchi *et al.* 2016). Treating Mesozoic bone that has experienced diagenesis, low latitudes, and permineralisation identically to more recent, less altered bone is ill-advised, and any work on such samples should employ these fundamental methods before attempting to sequence peptides that might not be present, ancient, or endogenous.

Fossil bone has fairly significant concentrations of recent organics (e.g., L-amino acids, DNA, and non-radiocarbon dead organic C), even when buried and often in comparison to the immediate environment. Fossil bone likely provides an ideal, nutrient-rich (e.g., phosphate, iron) open system microbial habitat inside vascular canals capable of moisture retention.

The absence of evidence for endogenous proteins and the presence of a diverse, microbial community urge caution regarding claims of dinosaur bone ‘soft tissues’. Microbes can colonise bones while buried, likely traveling via groundwater. Our results support the hypothesis that at least some ‘soft tissue’ structures derived from demineralised fossil bones represent biofilms. We suggest that unless in an inaccessible form (e.g., kerogen, depending on microbial metabolic ability) or matrix (e.g., well-cemented concretion), endogenous dinosaur organics that survive prior taphonomic processes (e.g., diagenesis) may be subject to subsequent microbial metabolic recycling.

The study of fossil organics must consider potential microbial presence throughout a specimen’s taphonomic history, from early to late. Microbial communities interact with fossils immediately following death and after burial, but prior to diagenesis. Microbes are known to utilize bone and tooth proteins (Child *et al.* 1993) and fossil evidence of early fungal colonisation has even been detected (Owocki *et al.* 2016). There is also recent microbial colonisation of fossil bone as it nears the surface in the late stages of the taphonomic process. Furthermore, given that microbes can inhabit the crust kilometres below the surface, it might be predicted that bone remains a biologically active habitat even when buried hundreds of meters deep for millions of years. The extensive potential for microbial contamination and metabolic consumption makes verifying claims of Mesozoic bone protein extremely challenging.

## Acknowledgements

Many thanks to Wei Wang and Jessica Wiggins (Princeton University, Genomics Core Facility) for 16S rRNA amplicon sequencing, Paul Monaghan (University of Bristol) for assistance in preparing samples for radiocarbon AMS, Sheila Taylor (University of York) for assistance in preparing the samples for HPLC, Kirsty High (University of York) for provision of the comparator sheep bone sample, Kentaro Chiba (University of Toronto) for assistance in the quarry of BB180 and for taking photographs, the Royal Tyrrell Museum of Palaeontology for accessioning the fossils collected in Dinosaur Provincial Park and assisting in the paperwork allowing for export and study, Mark Norell (American Museum of Natural History) for supplying a supplemental bone sample for comparison, and Adam Maloof (Princeton University), Jasmina Wiemann (Yale University), and Michael Buckley (University of Manchester) for helpful discussion. *Gallus gallus domesticus* (Public Domain Dedication 1.0, https://creativecommons.org/publicdomain/zero/1.0/legalcode), Odontapsidae (Dmitry Bogdanov, vectorised by T. Michael Keesey, Creative Commons Attribution 3.0 Unported, https://creativecommons.org/licenses/by/3.0/legalcode, CC BY 3.0), and *Centrosaurus apertus* (credit: Andrew A. Farke, Creative Commons Attribution 3.0 Unported, https://creativecommons.org/licenses/by/3.0/legalcode, CC BY 3.0, modified in Fig. 5) silhouettes were obtained from phylopic.org.

